# Research of the mechanism on miRNA193 in exosomes promotes cisplatin resistance in esophageal cancer cells

**DOI:** 10.1101/830240

**Authors:** Shifeng Shi, Xin Huang, Xiao Ma, Xiaoyan Zhu, Qinxian Zhang

## Abstract

**Purpose:** Chemotherapy resistance of esophageal cancer is a key factor affecting the postoperative treatment of esophageal cancer. Among the media that transmit signals between cells, the exosomes secreted by tumor cells mediate information transmission between tumor cells, which can make sensitive cells obtain resistance. Although some cellular exosomes play an important role in tumor’s acquired drug resistance, the related action mechanism is still not explored specifically.

**Methods:** To elucidate this process, we constructed a cisplatin-resistant esophageal cancer cell line, and proved that exosomes conferring cellular resistance in esophageal cancer can promote cisplatin resistance in sensitive cells. Through high-throughput sequencing analysis of the exosome and of cells after stimulation by exosomes, we determined that the miRNA193 in exosomes conferring cellular resistance played a key role in sensitive cells acquiring resistance to cisplatin. In vitro experiments showed that miRNA193 can regulate the cell cycle of esophageal cancer cells and inhibit apoptosis, so that sensitive cells can acquire resistance to cisplatin. An in vivo experiment proved that miRNA193 can promote tumor proliferation through the exosomes, and provide sensitive cells with slight resistance to cisplatin.

**Results:** Small RNA sequencing of exosomes showed that exosomes in drug-resistant cells have 189 up-regulated and 304 down-regulated miRNAs; transcriptome results showed that drug-resistant cells treated with drug-resistant cellular exosomes have 3446 high-expression and 1709 low-expression genes; correlation analysis showed that drug-resistant cellular exosomes mainly affect the drug resistance of sensitive cells through paths such as cytokine–cytokine receptor interaction, and the VEGF and Jak-STAT signaling pathways; miRNA193, one of the high-expression miRNAs in drug-resistant cellular exosomes, can promote drug resistance by removing cisplatin’s inhibition of the cell cycle of sensitive cells.

**Conclusion:** Sensitive cells can become resistant to cisplatin through acquired drug-resistant cellular exosomes, and miRNA193 can make tumor cells acquire cisplatin resistance by regulating the cell cycle.

## Introduction

Esophageal cancer is the eighth most common tumor in the world. Esophageal cancer patients in China account for more than half of the total number of esophageal cancer patients in the world, and here the mortalities of both male and female patients are the highest.^1,2^ The occurrence of esophageal cancer is affected by multiple factors, including genetics, living environment, bad habits (such as smoking and drinking) and others.^1^ It is not easy to detect esophageal cancer at an early stage, and esophageal cancer in the middle and advanced stages is generally treated with chemotherapy and radiation. As a broad-spectrum antitumor drug, cisplatin mainly causes DNA damage in tumor cells, and is a common chemotherapeutic drug used to treat esophageal cancer.^3,4^ However, drug resistance generated by esophageal cancer cells is a decisive factor that affects the chemotherapeutic effects.

Exosomes, nanoscale vesicles with lipid bimolecular films, are a type of extracellular vesicles (EV) generated and released by most cells.^5^ Exosomes occur in all body fluids,^5–8^ and due to their potential effects as messengers between cells and as new non-invasive tumor biomarkers,^9,10^ exosomes have attracted broad attention in recent years. The exosomes secreted by tumor cells play a main role in transmitting information from tumor cells to other malignant or normal cells,^6,11,12^ and can be regarded as the medium to transfer information.

The exosomes secreted by tumor cells mediate information transfer between tumor cells (drug-resistant cells and sensitive cells), which can make sensitive cells obtain drug resistance.^6,13^ Wei et al found that by treating the tamoxifen-sensitive breast cancer cell line MCF-7 with the exosome secreted by chemotherapy-resistant cell line MCF-7^TamR^, it can acquire drug resistance, because the miR-221/222 in the exosome secreted by the drug-resistant cell line inhibited the expression of the estrogen receptor α target gene (ERα).^14–16^ miR-222 can make the chemotherapy-resistant breast cancer cells regain their sensitivity to adriamycin.^17^ In our research, we found that the exosome of the cisplatin-resistant esophageal cancer cell line can induce sensitive cells to become resistant to cisplatin. By combining high-throughput sequencing technology with later cytobiological verification, we found that this phenomenon may be related to the cell cycle.

## Materials and methods

TE-1 cells were provided by the Experiment Center, School of Basic Medical Sciences, Zhengzhou University, and were bought from the Cell Resource Center, Shanghai Institutes for Biological Sciences. The cells were cultured in RPMI-1640 medium containing 10% exosome-depleted fetal calf serum (Gibco; Thermo Fisher Scientific, Inc., Waltham, MA, USA) and 100U/ml penicillin (Invitrogen; Thermo Fisher Scientific). All cell cultures were maintained in an incubator containing 5% CO_2_ at 37°C.

Following the method of Jiang, cisplatin (cis-diaminedichloroplatinum, DDP; Sigma, St. Louis, MO, USA) was added to the normally cultured TE-1 cells in the medium.^18^ First, the CCK-8 kit (Genecopeia, Rockville, MD, USA) was used to test the median inhibitory concentration (IC_50_) of sensitive cell TE-1 to DDP as 0.25 μM; then, the medium containing 3.3 μM DDP was used for a 24-h culture; after that, the medium containing DDP at median inhibitory concentration was used to incubate seven days until cell growth became stable and continuous passage for 3 times; after measurement of IC_50_, the medium containing 3.3 □M DDP was used for culture and shock for 24 h; then the medium was replaced with medium with a gradually increasing concentration of DDP. When the cell displayed stable growth in the medium containing 3.3 μM DDP and had continuous passage for 3 times, we measured the IC_50_ of DDP as 8.2 μM, and named the drug-resistant cell line as TE-1/DDP. TE-1/DDP was cultured in the RPMI-1640 medium containing 10% exosome-depleted fetal calf serum (Gibco; Thermo Fisher Scientific), 2 μM DDP and 100U/ml penicillin (Invitrogen; Thermo Fisher Scientific) to maintain resistance to cisplatin.

### Exosome separation and identification

The exosomes from the TE-1 and TE-1/DDP cell supernatants are referred to as TE-1/exo and TE-1/DDP/exo, respectively. The cell supernatant was centrifuged at 1,000 × *g* (10 minutes), 10,000 × *g* (30 minutes at 4°C) and 100,000 × *g* (120 minutes at 4°C) by using a Beckman ultracentrifuge (AvantiJ-30I; Beckman Coulter, Inc., Brea, CA, USA). The final sediment was resuspended in the ExoQuick-TC exosome precipitation solution (System Biosciences, Mountain View, CA, USA) at 4°C overnight, and centrifuged at 1,500 × *g* (30 minutes at 4°C). The extracted sediment was diluted in 200 μl phosphate-buffered saline (PBS) and stored at −80°C. A JEM-2100 transmission electron microscope (TEM; JEOL, Ltd., Tokyo, Japan) was used to observe the exosomes. Western blotting was used to measure the cell surface marker CD63.^19 20^

### Test of cell vitality

The day before experiments, pancreatin was used to digest the cells, and a cytometer (Countstar; Countstar BioTech, Shanghai, China) was used to count the cell suspension. Cells were inoculated with a density of 3500 cells/well in the 96-well cell culture plate, and after culture for 24 h, cells were divided into groups that were processed differently. In accordance with the manufacturer’s operating instructions, the CCk-8 kit was used to test the cell vitality.

### Small RNA high-throughput sequencing analysis

TRIzol was used to extract total RNA from the sample, and AGE (agarose gel electrophoresis) was used to cut and select 18–30 nucleotide (nt) segments. The 3’ joint and 5’ joint were connected, respectively; then, reverse transcription and PCR amplification were conducted on the small RNAs with joints at both ends; finally, AGE was used to recover and purify a ladder of around 140 base pairs (bp), and library construction was completed. Agilent 2100 (Agilent, Santa Clara, CA, USA) and qPCR were used for quality control of the constructed library, and computer-controlled sequencing was conducted.

For the small RNA sequencing data, first of all, we used a dynamic programming algorithm to filter out low-quality reads that included polyA, and sequences with reads smaller than 16 nt or larger than 35 nt were all eliminated. After filtering, the quality and length distribution of the sequencing data were summarized. Then, standard bioinformatics analysis of the small RNA sequences was conducted. We compared the remaining sequences with the NCBI non-coding RNA database (http://blast.ncbi.nlm.nih.gov/) and the Rfam database (ftp://sanger.ac.uk/pub/databases/Rfam/) to separate rRNA, tRNA, snRNA and other ncRNA sequences. Then, miRNA identification was conducted by comparing the sequence with the known mature miRNA and miRNA precursors in miRBase 21.0 (http://www.mirbase.org/) to identify the known miRNAs. In the meantime, miRDeep (http://deepbase.sysu.edu.cn/miRDeep.php) was used to analyze many unannotated sequences that could not be matched with the above databases to predict new miRNA candidates. Then, the RNAfold software (http://rna.tbi.univie.ac.at/cgi-bin/RNAfold.cgi) was used to analyze their hairpin structure.

The total miRNA reads were standardized to select the miRNAs with differential expressions between the two libraries. Per million (TPM) was used to standardize the miRNA counts. Old known and newly known miRNA expression levels between the two libraries (TE-1/exo was used as control group) were compared to find the differential expressions of the miRNA. Here, the selection method was based on the method of Audic and Claverie.^21^ The IDEG6 software was used to analyze differential expressions of miRNA.^22^

The exosomes in TE-1/DDP medium were collected by ultracentrifugation. The collected TE-1/DDP/exo were added to the medium to culture cisplatin-sensitive TE-1 cells; the cells were collected after culture for 24 h and are referred to as TE-1-Exo. After collecting the TE-1 and TE-1-Exo cells, the TRIzol reagent was used to extract total RNA, and RNA employed NanoDrop 2000 (Thermo Scientific) was applied to measure the concentration of each sample. Agilent2200 (Agilent) was used to evaluate the integrity of RNA. The RNAs with RIN (RNA integrity number) values of 8.3–9.1 were used for sequencing.

For the original sequencing data of the transcriptome, we adopted the Perl script for data filtering and screening of effective data. All downstream analyses were based on high-quality effective data. Bowtie v2.2.3 was used to build the reference genome index, and TopHat v2.0.12 was used to compare the paired-end clean reads with the reference genome. The edgeR R package was used to identify the genes with differential expressions between the TE-1-Exo group and TE-1 group. All data were normalized to generate unsupervised clustering analysis heat maps.

A gene ontology (GO) enrichment analysis was conducted on the differential genes obtained through screening, and when *p* <0.05, the GO terminology is regarded as significantly enriched. The Kyoto Encyclopedia of Genes and Genomes (KEGG, https://www.genome.jp/kegg/kegg2.html) was used for gene enrichment of differentially expressed genes. The protein–protein interaction (PPI) analysis of differentially expressed genes was based on the STRING database v10.0 (https://string-db.org/cgi/input.pl), and the threshold of STRING was set at 500.

### Quantitative real-time polymerase chain reaction

According to the manufacturer’s recommendation, we used the Trizol reagent to separate the total RNA. The Nanodrop 2000 spectrophotometer (Thermo Scientific, Wilmington, DE, USA) was used to evaluate the quality and purity of RNA. All-in-One™ First-Strand cDNA Synthesis Kits (Thermo Scientific) were used to reverse transcript the RNA sample (1 μg) to cDNA; All-in-One qPCR mix (Thermo Scientific) was used to measure the expression level of mRNA according to the SYBRGreenI method. All-in-One™ miRNA qRT-PCR Detection Kit (Genecopeia, Rockville, MD, USA) was used in the reverse transcription and test of miRNA to analyze the relative expression of miRNA in the two libraries.

The PCR measurement was conducted with a fluorescent quantitative PCR instrument (Bio-Rad CFX96, Hercules, CA, USA). The primer sequences are listed in Table S1. The thermal program included the following melting curve steps: 1 minute at 95℃, 10 cycles and 40 cycles at 95℃, 30 s at 60℃ and a gradual increase from 65℃ to 95℃ at 0.5°C s^−1^; the data were collected every 6 s. Each miRNA and gene expression through qRT-PCR ration was represented as *n* = 1 to complete for three times, and the U6sniRNA and β-actin genes were respectively used as internal controls. The Ct value is presented as the average value of three independent repetitions; the relative expression level was calculated with the □□Ct method. The standard error of the mean for the repetitions was also calculated.

### Western blot

Cold lysis buffer containing 50 mmol/l Tris–HCl (pH 7.4) was used to lyse the obtained cells. The DC protein assay kit (Bio-Rad Laboratories) was used to test the protein content, and a western blot was conducted according to the scheme of Hnasko’s protocol.^23^ Antibodies TFAP2C, TP53, P21, EGFR, Cylin D1, Bcl2, Bax, cCaspase-3 and β-actin were all bought from Abcam (Cambridge, UK). The enhanced chemiluminiscence solution (Life Sciences; Thermo Fisher Scientific) was used to visualize proteins.

### Dual luciferase reporter

The 3’ untranslated region in the TFAP2C of the miR-193a target gene and its mutant were cloned into the pEZX-MT01 carrier (Genecopeia, Rockville, MD, USA). The miR-193a or miR-NC overexpressed plasmids (Ribobio, Guangzhou, China) and the firefly luciferase reporter carrier were co-transfected to human embryonic kidney (HEK293) cells. The dual luciferase assay kit (Promega, Madison, WI, USA) and Glomax 96 Microplate Spectrophotometer (Promega, Madison, WI, USA) were used to measure the firefly and sea pansy luciferase activity 48 h after co-transfection.

### Test of cell cycle

The PBS solution was used to process around 1 × 10^6^ cells to single-cell suspension, and it was prepared according to the manufacturing plan for the annexin V-FITC apoptosis assay kit (Beyotime Biotechnology, Haimen, Jiangsu, China). Then, the FACScan system (BD Biosciences, San Jose, CA, USA) was used to analyze the apoptosis rate.

The PBS solution was used to process around 1 × 10^6^ cells to single-cell suspension twice, which was fixed with 70°C ice alcohol for 24 h. Then, after washing and centrifugation with PBS, the propidium iodide solution (containing 100 mg/l RNaseA) was added, and it was incubated at room temperature away from the light. Then, the FACScan system (BD Biosciences, San Jose, CA, USA) was used to measure the cell cycle distribution, and it was analyzed with Multicycle for Windows software (Beckman Coulter, California, USA).

### Nude mice bearing tumor experiment

The six-weeks-old female mice were bought from Beijing Vital River Technology Co., Ltd., and they were kept in the animal facilities of the Medical School, Zhengzhou University, before being used in the experiments. Subcutaneous inoculation of tumor cells was conducted at the upper limbs on both sides of the mouse spine, and the inoculation dosage was 5 × 10^5^ cells/mouse. The TE-1 cells were inoculated on the right side, and the TE-1/DDP, TE-1/miR193 and TE-1/mimic-NC cells were inoculated on the left side. After the tumor grew to a size of 2 mm^3^, nude mice inoculated with the same cells were randomly divided into two groups (each group consisted of *n* = 4 mice). Then, cisplatin or normal saline (100 mg/kg/day) was injected into the body of the mice by intraperitoneal injection every day, and this lasted for seven days. On the seventh, tenth, fourteenth, sixteenth, eighteenth and twenty-first days, the tumor size was measured with calipers. The tumor volume was estimated according to the formula of (*L* × *S*^2^)/2 (where *L* is the longest diameter and *S* is the shortest diameter). The mice were killed, the tumors were collected and weighed with an electronic scale (ME104, Mettler Toledo, Shanghai, China). The experiment was approved by the Ethics Committee of Medical School, Zhengzhou University.

### Statistical analysis of data

SPSS software (version 22.0; IBM Corp., Armonk, NY, USA) was used for statistical analysis. Student’s *t*-test was used for paired comparison, one-way analysis of variance was conducted, and then, the Student Newman–Keuls test was used for multiple comparisons between groups. All experiments were conducted three times, and the results are presented as mean values ± standard deviation. When *p* < 0.05, the difference is regarded as statistically significant.

### Results

### Drug-resistant cell exosome makes sensitive cells acquire resistance to cisplatin

In order to evaluate the tolerance of the constructed cisplatin-resistant esophageal cancer cell line TE-1/DDP to cisplatin, TE-1 and TE-1/DDP were cultured in the cisplatin medium containing IC_50_TE-1 (2.5 μM), and the vitality of the TE-1/DDP cell line was significantly higher than that of the TE-1 cell line (*p*<0.001) (Fig. 1A).

**Figure 1.**
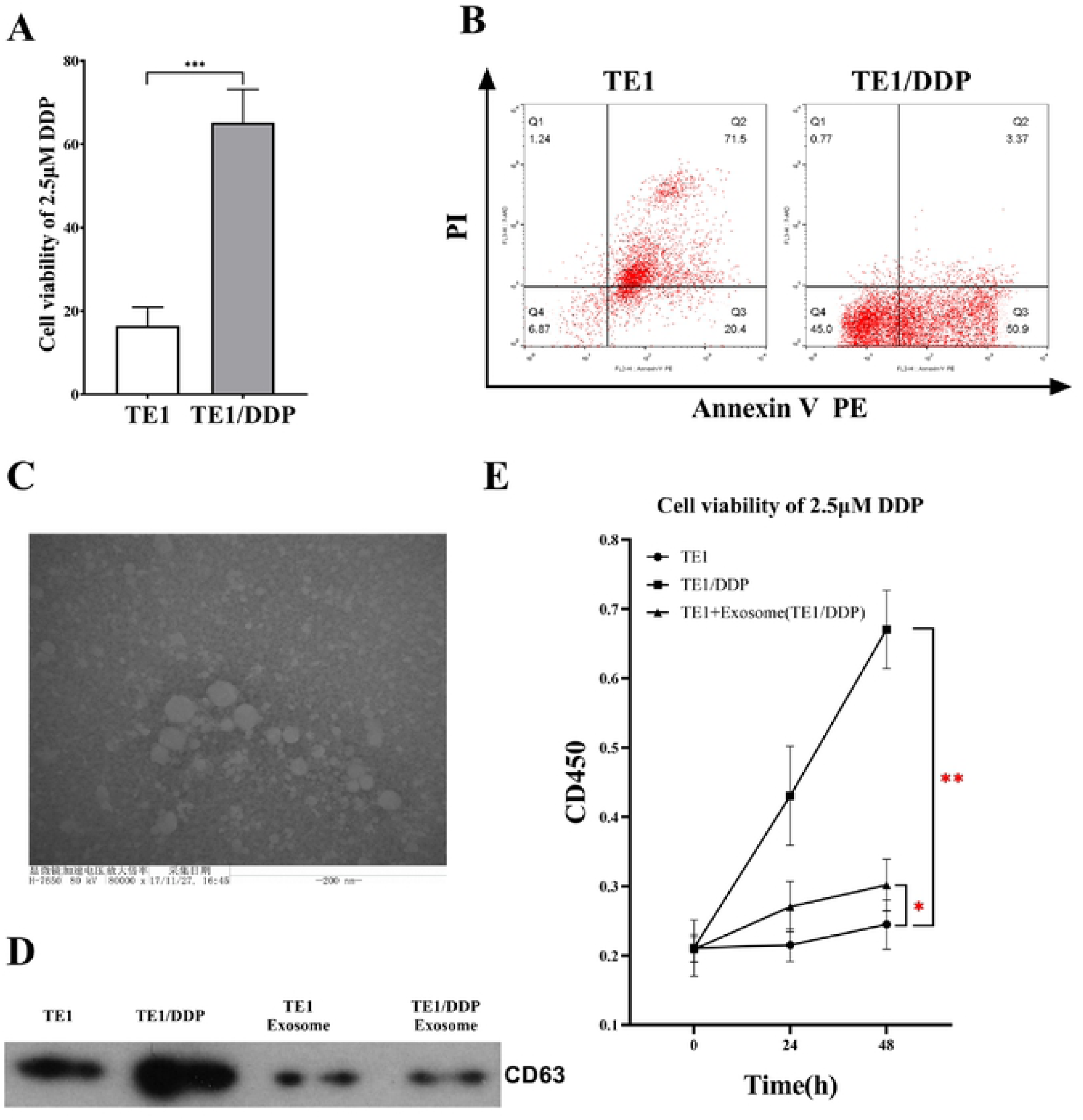
Identification of drug-resistant cell lines and exosomes. A: Detection of cell viability of original cell line (TE-1) and drug-resistant cell line (TE1/DDP) under 2.5 μM DDP conditions. B: Apoptosis detection in original cell line (TE-1) and drug-resistant cell line (TE1/DDP) under 2.5 μM DDP. C: Scanning electron microscopy to identify exosomes. D: Western blot analysis of good exosome membrane protein marker CD63. E: CCK-8 detects cell viability showing that drug-resistant exosomes (TE1/DDP-Exo) *promote* DDP resistance in TE1 cells.

The flow cytometer test results showed that 24 h after being processed with 2.5 μM cisplatin, apoptosis was almost complete in the TE-1 cell line and the apoptosis rate was 98.4 ± 1.2%, while the cisplatin-resistant esophageal cancer cell line TE-1/DDP was only partially apoptotic, and the apoptosis rate was 52.8 ± 15.9%. Upon comparison, we found the difference was statistically significant (*p* < 0.001) (Fig. 1B).

Under the transmission electron microscope the esophageal cancer cell exosome presented the classical round or oval cup-shaped form, and its size was around 50–200 nm (Fig. 1C). Western blotting was used to detect the exosome’s marker protein molecule CD63. As a result, the CD63 protein was detected in the TE-1 cell and the TE-1/DDP cell, as well as in the extracted exosomes of the TE-1/DDP cell and the TE-1 cell (Fig. 1D).

The CCK-8 kit was used to test the cell vitality, and we identified the influence of TE-1/DDP exosome on the tolerance of TE-1 cells for cisplatin. The results showed that under the IC_50_ TE-1 (2.5 μM) cisplatin pressure, the vitality of the TE-1/DDP cell line was significantly higher than that of the TE-1 cell line (*p* < 0.001); after adding the TE-1/DDP exosome, the TE-1 cells’ tolerance of DDP was significantly increased (*p* < 0.05) (Fig. 1E).

### Correlation analysis between small RNA-Seq and RNA-Seq

The Illumina HiSeq 2500-high throughput technology was used to sequence the small RNA libraries of two exosomes (TE-1 and TE/DDP). At the beginning, the original reads of 12604198 and 17480281 were generated. After removing low-quality reads, the adapter sequences and reads shorter than 18 nt, clean reads of in total 7879335 and 12960223 were obtained. The sequence length distributions of the two libraries presented significant differences; most sequences were between 21 and 23 nt (Fig. 2A). Next, we mapped all clean reads to miRBase (v.20) to annotate the known miRNAs in each library. The reads of TE-1/exo and TE-1/DDP/exo were annotated to obtain 632 and 712 miRNAs, respectively. However, in this research, we decided to use a stricter threshold (> 5 TPM cutoff value), so that we could focus on representative miRNA. As a result, 493 different miRNAs from the TE-1 group and the TE-1/DDP group were used for further analysis. Compared to the TE-1/exo group, 189 were up-regulated miRNAs, and 304 were down-regulated miRNAs, including the hierarchical clustering of expression level (Fig. 2B).

**Figure 2.**
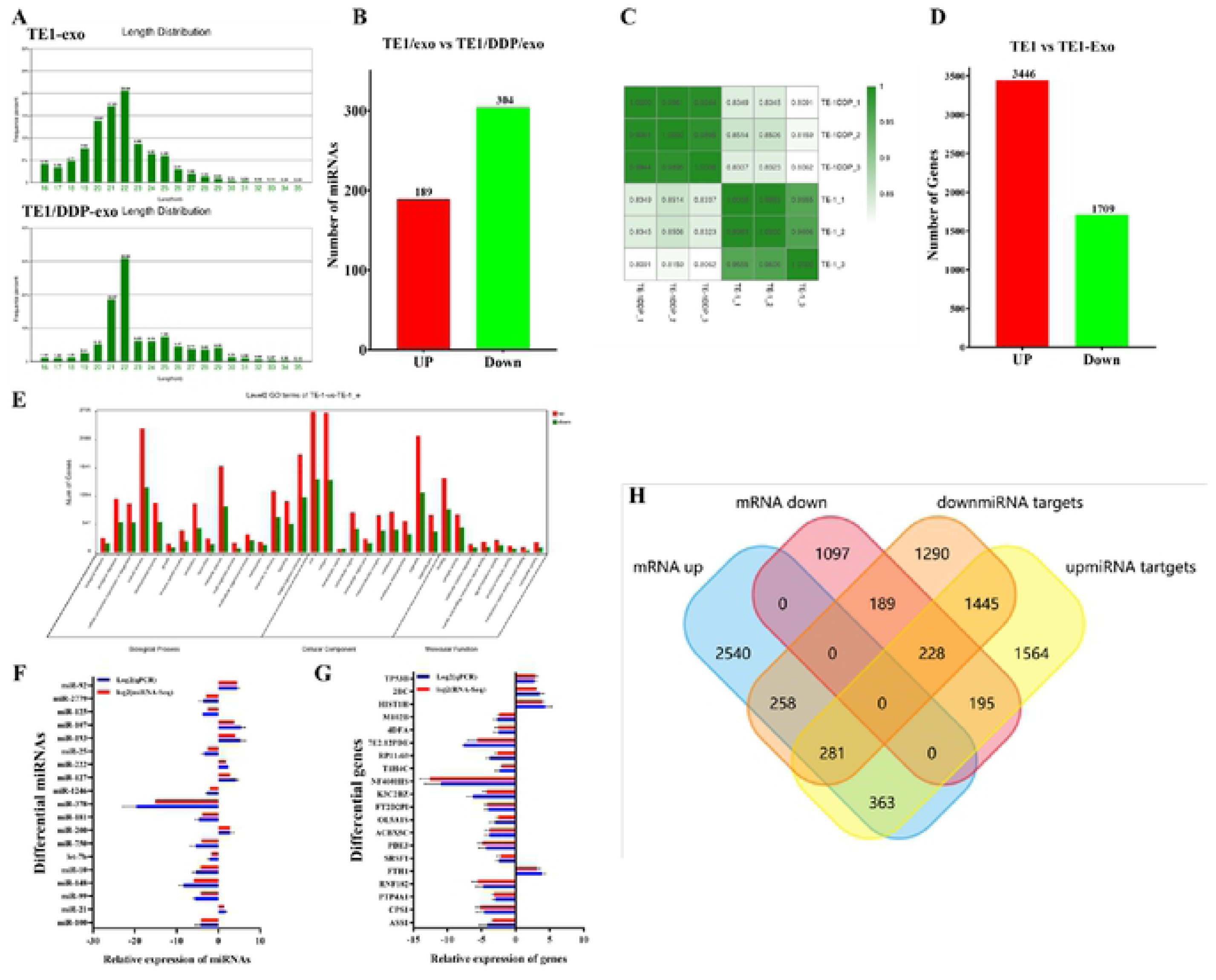
Analysis and identification of next-generation sequencing results. A: miRNA length distribution in different exosomes samples. B: Difference in miRNA expression levels in exogenous samples from different sources. C: Sample correlation analysis between sample sequencing data sets. D: Analysis of difference in gene expression between cells after drug-resistant exosome stimulation. E: Differential gene Go analysis. F, G: RT-qPCR validation of the significantly different expression of miRNAs and genes. H: Wayne diagram analysis shows the intersection of miRNA target genes and differential genes.

After sequencing, the adapter sequences, fuzzy reads and low-quality reads were eliminated, and around 55 × 10^5^ to 110 × 10^5^ clean reads were generated for each sample. Compared to the reference sequence of genome complex GRCh37/hg19, 92% of all reads were uniquely positioned in the human genome. The mapped reads were used to estimate the normalized transcriptional level as FPKM. Correlation matrix analysis and principal component analysis (PCA) were employed to estimate the clustering properties of these samples. The samples in each group were aggregated, and in this way displayed great repeatability and correlation (Fig. 2C).

In total, 5155 differential expression genes were determined as being related to the effects of TE-1/DDP/exo. Of these, 3446 genes had up-regulated expression, and 1709 genes had down-regulated expression (Fig. 2D). In order to further analyze the function of these differential genes, we conducted GO analysis based on the GO terminologies (Fig. 2E). We identified 16 overrepresented GO terminologies related to the biological process (*p* < 0.05). These terminologies included genes that participated in cell proliferation, cell adhesion, angiogenesis regulation and metabolic processes. Eight overrepresented GO terminologies were related to the molecular function (*p* < 0.05). These genes belonged to the genes with nucleic acid binding transcription factor activity, protein binding transcription factor activity, signal transduction activity and structural analysis activity. Ten GO terminologies with significant difference (*p* < 0.05) were related to the components of cells, and they mainly belonged to the extracellular components and organelles (Fig. 2E).

In order to verify the expression level of miRNAs with differential expression (TOP24) in the sequencing data of TE-1/exo and TE-1/DDP/exo, we used miRNA fluorescent quantitative PCR to measure the expression levels of miRNAs with significant difference (Fig. 2F). The same method was employed to measure the expression level of Top20 mRNA with significant difference in the transcriptome (Fig. 2G). The results showed that the expression levels of some of the miRNAs and mRNA with significant differences screened by us in the separate exosomes or cells were consistent with the expression results of the sequencing data.

The bioinformatics software FunRich3.1(http://www.funrich.org) was used to conduct correlation analysis of the up-regulated mRNAs and down-regulated miRNA target genes (Fig. 2H). By searching the GO and KEGG analyses, we found that the gene change caused by stimulation of the drug-resistant cellular exosomes mainly affected the cytokine–cytokine receptor interaction, and the Jak-STAT signaling, VEGF signaling and oxidative phosphorylation pathways (Table 1).

**Table 1.** Go and KEGG analysis of Top10 differential gene pathway richment

| Go and KEGG analysis of Top10 differential gene pathway richment |  |  |  |  |  |
| --- | --- | --- | --- | --- | --- |
| NO | Pathway | Gene number | All (55772) | Pvalue | Qvalue |
| 1 | Lysosome | 53 (3.09%) | 123 (1.82%) | 0.000013 | 0.001255 |
| 2 | Oxidative phosphorylation | 52 (3.03%) | 132 (1.96%) | 0.000264 | 0.019159 |
| 3 | AGE-RAGE signaling pathway in diabetic complications | 40 (2.33%) | 100 (1.48%) | 0.000918 | 0.045512 |
| 4 | Transcriptional misregulation in cancers | 63 (3.67%) | 174 (2.58%) | 0.000942 | 0.045512 |
| 5 | JAK-STAT signaling pathway | 70 (4.08%) | 200 (2.96%) | 0.001459 | 0.060449 |
| 6 | VEGF signaling pathway | 26 (1.51%) | 60 (0.89%) | 0.001851 | 0.067112 |
| 7 | Pathways in cancer | 122 (7.11%) | 388 (5.75%) | 0.00367 | 0.106421 |
| 8 | Rap1 signaling pathway | 70 (4.08%) | 214 (3.17%) | 0.009453 | 0.210876 |
| 9 | Protein digestion and absorption | 32 (1.86%) | 88 (1.3%) | 0.014687 | 0.266209 |
| 10 | ECM-receptor interaction | 29 (1.69%) | 80 (1.19%) | 0.020452 | 0.327602 |

### miRNA193 increases resistance to cisplatin by regulating TFAP2C

The small RNA-seq results showed that miRNA-193 is a high-expression miRNA in TE-

1/DDP/exo. Similarly, by analyzing the transcriptome data, we found that in TE-1/exo, the expression level of TFAP2C was significantly lower than that of the TE-1 cells. A fluorescent quantitative PCR experiment was conducted to measure the miRNA193 and TFAP2C in different samples, which showed similar differences (Fig. 3A). TargentScan-Human V7.0 (http://www.targetscan.org) was used to predict that miRNA-193 and TFAP2C had a targeting sequence relation (Fig. 3B). The test results of the dual luciferase reporter showed that miR193 was only combined with correct TFAP2C gene 3’-UTR area, indicating that TFAP2C was the target gene of miR193 (Fig. 3C).

**Figure 3.**
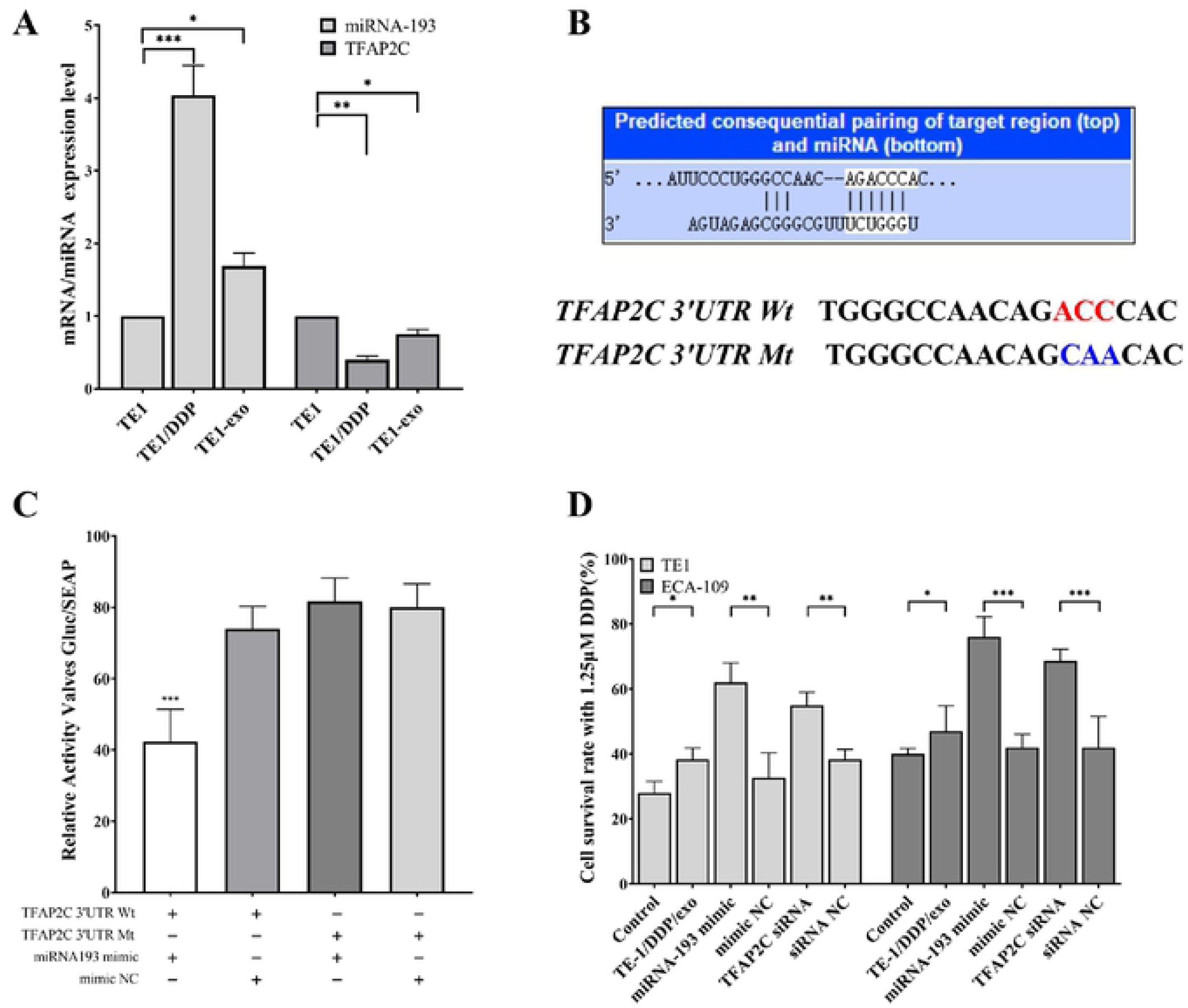
Verification that TFAP2C is the target gene of miRNA193. A: qPCR detection of differential expression of miRNA193 and TFAP2C in different cells. B: TargentScan 7.2 predicts the targeting relationship between TAFAP2C and miRNA193, and its dual luciferase experimental mutation site. C: Dual luciferase assay for targeted binding of miRNA193 to TFAP2C. D: Changes in resistance to cisplatin in esophageal cancer cells after addition of TE1/DDP-Exo, transfection of miRNA193 minic or TFAP2C siRNA.

We added TE-1/DDP/exo to the esophageal cancer cell TE-1 and breast cancer cell ECA-109, which were sensitive to cisplatin. In the meantime, TE-1, ECA-109, the miRNA193 mimic, and TFAP2C were transfected to measure the cell vitality when 1.25 □M DDP was added to TE-1 and ECA-109 cells. In this process, normal TE-1 and ECA-109 were used as the control group. The results showed that the cisplatin resistance of cells was increased when TE-1/DDP/exo was added. In TE-1 and ECA-109 cells, the overexpressed miRNA193 or down-regulated TFAP2C enabled the cells to acquire tolerance to cisplatin (Fig. 3D).

We wished to prove the hypothesis that exosomes or miRNA19 could cause drug resistance by an experiment with nude mice bearing tumors. Cell inoculation was conducted in the different groups as shown in the diagram (Fig. 4A). Among them, for Group A and Group D, TE-1/DDP was injected into the left underarm of nude mice (L), and TE-1 was inoculated in the right underarm (R). For Group B and Group E, TE-1/miR193 mimic was inoculated into the left underarm of nude mice, and TE-1 was inoculated in the right underarm. For Group C and Group F, TE-1 was inoculated in both underarms of nude mice. Three weeks after cell inoculation, Groups A–C were injected with normal saline, and they were used as the control group. Groups D, E and F were injected with cisplatin through intraperitoneal injection.

**Figure 4.**
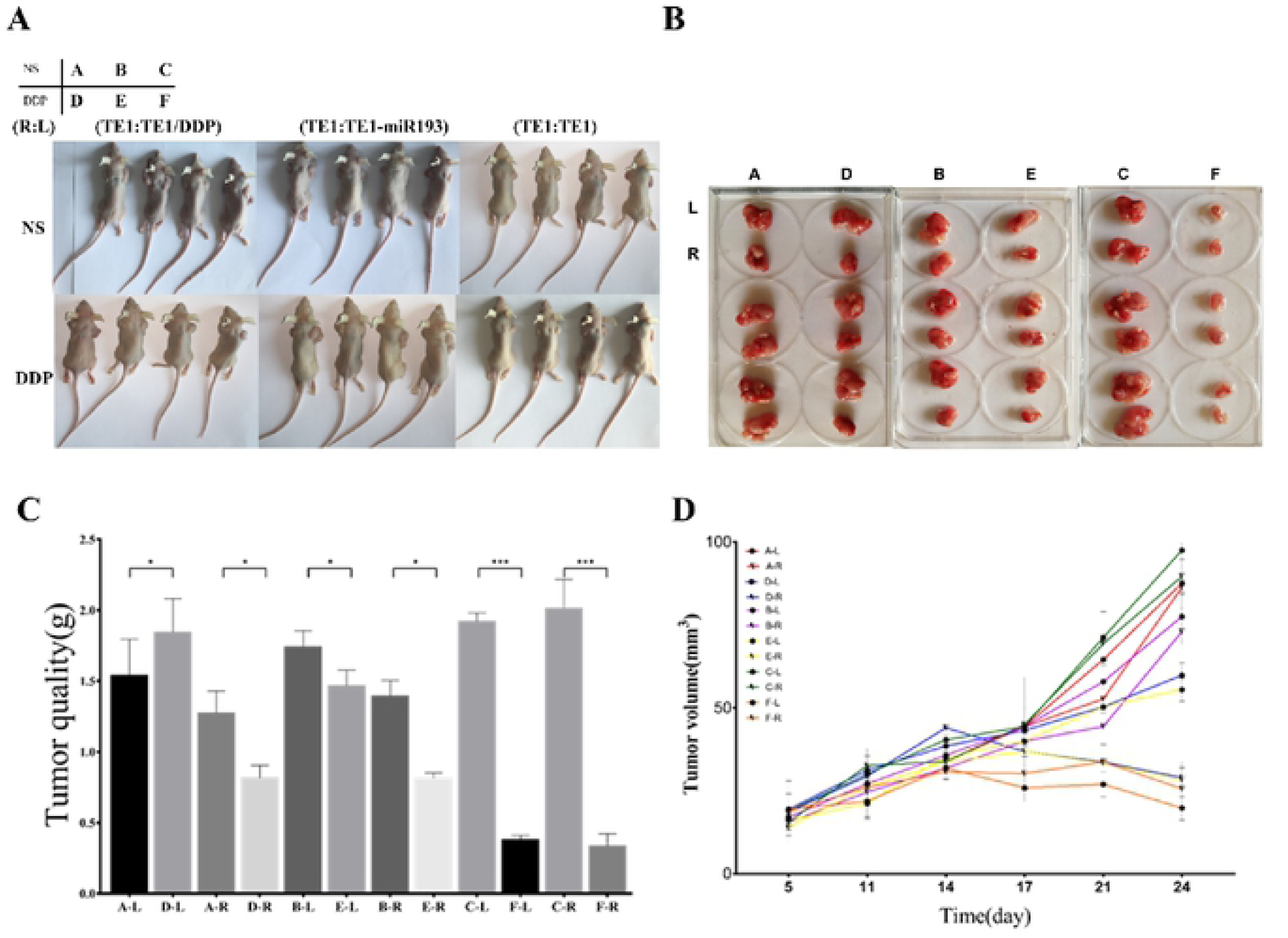
In vitro experiments demonstrate that both drug-resistant exosomes and miRNA193 increase the sensitivity of sensitive cells to cisplatin. A, B: nude mice and postoperative tumor photos. C: Comparison of tumor size in different groups of nude mice. D: Tumor growth curves in different groups and different locations.

The results of the experiment with nude mice bearing tumors showed that under intervention with cisplatin, the size of the formed tumor with TE-1/DDP was slightly smaller than that of the normal saline group (Group A-L compared with Group D-L). Under the same intervention, the size of the formed tumor in the TE-1 cell line was significantly smaller than that of the normal saline group (Group C compared with Group F). Under the intervention with cisplatin, the size of the formed tumor of TE-1/miRNA193 mimic was smaller than that of the normal control group (Group B-L compared with Group E-L). Under the intervention with cisplatin, sizes of the formed tumors of TE-1 cells in Group B and Group E were significantly larger than the tumor size of Group F (Fig. 4B–D). The above results prove that cisplatin can inhibit the growth of esophageal cancer tumor cell TE-1. The TE-1/DDP cells can significantly resist the anti-tumor effects of cisplatin, and overexpression of miR193 can promote the growth of esophageal cancer tumor. The exosomes secreted by the TE-1/DDP and TE-1/miRNA193-mimic cells can make sensitive TE-1 cells generate resistance to cisplatin when they circulate in the body fluid.

### High-expression miRNA193 or low-expression TFAP2C can remove cell cycle inhibition and inhibit apoptosis

We employed flow cytometry to test the influence of changes in TFAP2C and miR193 expression on the cell cycle. The proportion of G0/G1-stage cells in TE-1/TFAP2C-OE was significantly higher than that in the control group TE-1/OE-control. The proportion of G0/G1-stage cells in TE-1/miRNA mimic was significantly lower than that in TE-1/mimic-NC. In the medium containing 0.25 μM cisplatin, the proportion of G0/G1-stage cells of TE-1 cells was higher than the proportion of normally cultured TE-1 cells (Fig. 5A). Therefore, it can be seen that cisplatin can inhibit the cell cycle of esophageal cancer. The overexpression of miR193 can promote the division and proliferation of esophageal cancer cells, and the overexpression of TFAP2C can inhibit the division and proliferation of esophageal cancer cells.

**Figure 5.**
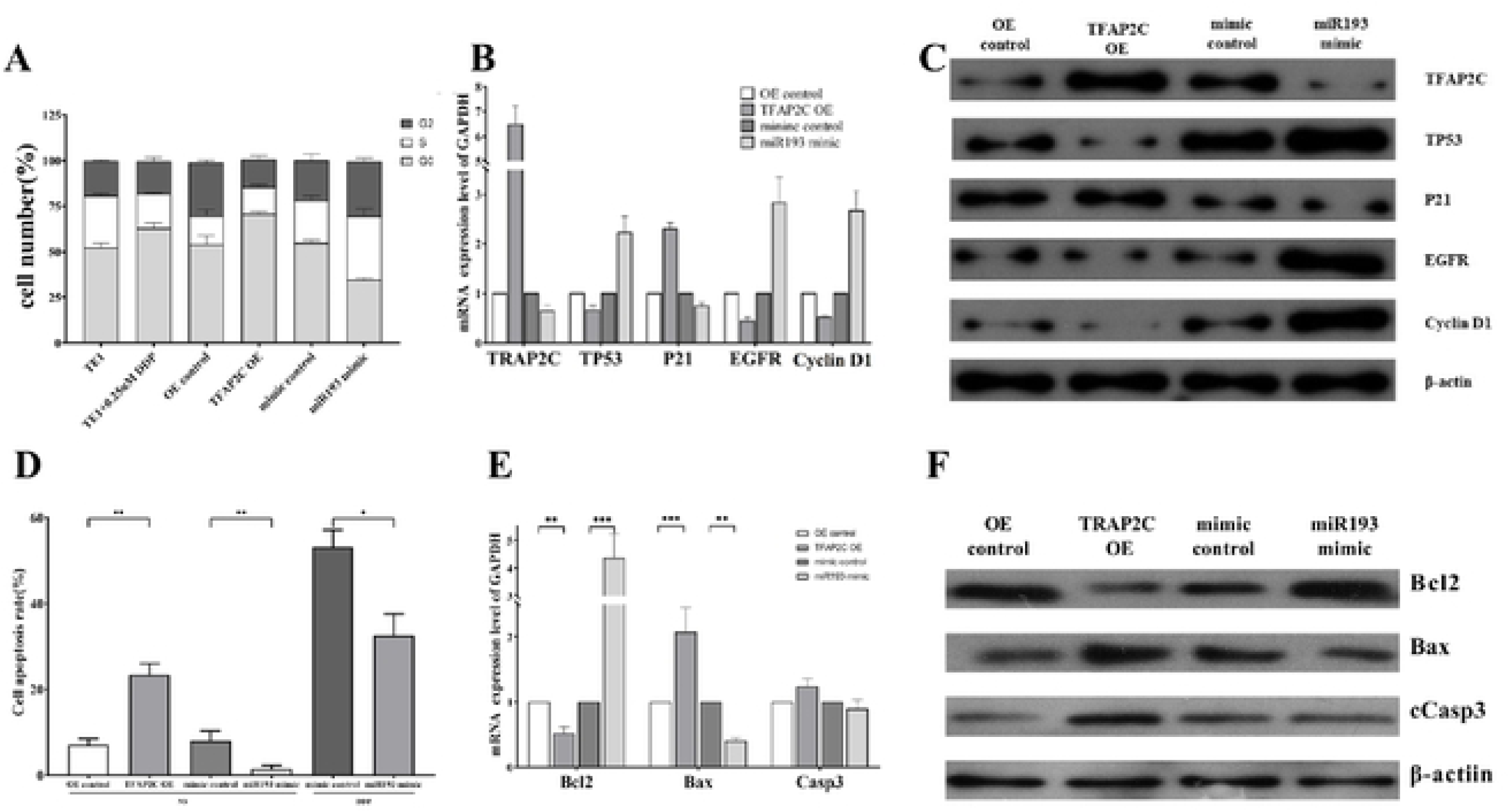
TFAP2C and mIRNA193 affect cell cycle and apoptosis. A: Effect of DDP, transfection of miRNA193 mimic or TFAP2C siRNA on TE1 cell cycle. B: qPCR detection of mRNA expression differences of cell cycle-related genes in differently treated cells. C: Western blot analysis of protein expression differences in cell cycle-related genes in differently treated cells. D: Apoptosis assay showing that overexpression of TFAP2C promotes apoptosis, and overexpression of miRNA193 inhibits cisplatin-induced apoptosis of esophageal cancer cells. E: qPCR detection of mRNA expression differences of apoptosis-related genes in differently treated cells. F: Western blot analysis of protein expression differences in apoptosis-related genes in differently treated cells.

We further adopted the RT-qPCR technology and western blot to test the influence of overexpressed TFAP2C or overexpressed miR193 in the TE-1 cell line on the expression of cell cycle-related genes. The results showed that compared to TE-1/OE-control, the relative mRNA expression level of TP53, EGFR and Cyclin D1 genes in TE-1/TFAP2C-OE significantly declined, but the relative expression level of P21 increased greatly. Compared to TE-1/mimic-NC, relative mRNA expression level of TP53, EGFR and Cyclin D1 genes in TE-1/miRNA193 mimic increased significantly, but relative mRNA expression level of P21 significantly decreased. The western blot results were similar to the fluorescent quantitative PCR results (Fig. 5B, C).

Flow cytometry results showed that compared to TE-1/OE-control, the apoptosis rate of TE-1/TFAP2C-OE significantly increased. Compared to TE-1/mimic-NC, the apoptosis rate of TE-1/miR193 mimic significantly declined. In the medium containing 0.25 μM cisplatin, the apoptosis rate of TE-1/miR193 mimic was significantly lower than that of TE-1/mimic-NC (Fig. 5D). These results showed that the overexpression of TFAP2C can promote the apoptosis of esophageal cancer cells, while the overexpression of miR193 can inhibit the apoptosis of esophageal cancer cells caused by cisplatin.

The RT-qPCR results showed that compared with TE/OE-control, the relative expression level of Bcl2 in TE-1/TFAP2C-OE significantly decreased, while the relative expression levels of Bax and Casp3 increased. Compared with TE-1/mimic-NC, the relative expression level of Bcl2 in TE-1/miR193 mimic increased, while the relative expression levels of Bax and Casp3 significantly declined. The western blot results were consistent with the qPCR results (Fig. 5E, F). Therefore, it can be seen that overexpression of TFAP2C can cause an increase of activated caspase-3, which will result in apoptosis. Overexpressed miRNA193 can inhibit TFAP2C, which generated the opposite result.

## Discussion

Drug resistance is a very severe problem that has not yet been solved. In tumor, most exosomes are the promoting factors of tumor development,^24–26^ as well as key components of the tumor microenvironment. Exosomes can be used as carriers for tumor signal transmission, and it is worth mentioning that, as the source of tumor, one of the negative clinical effects of exosomes is their ability to horizontally transfer drug resistance.^6,13,27^ Drug-resistant tumor cells can transfer drug resistance to sensitive cells through exosomes, thereby generating a new anti-tumor cell repository. For example, when exosomes were transferred from adriamycin- and docetaxel-tolerant breast cancer-resistant cell lines to the sensitive cell line, some miRNAs (miR-100, miR-222, miR-30a, and miR-17) would generate drug resistance.^17, 28–30^ In studies on prostatic cancer and docetaxel resistance, the exosomes secreted by drug-resistant cells can regulate the expression of target cell translocators and endow sensitive cells with docetaxel resistance.^31^ According to our research, levels of miRNA193 in esophageal cancer drug-resistant cell line TE-1/DDP and exosome TE-1/DDP/exo significantly increased. When TE-1/DDP/exo was added to the TE-1 cells, the miRNA193 in TE-1-Exo cells also increased to some degree, showing that the miRNA193 in exosome can be transferred to the target cells.

The activator 2 (TFAP2) family consists of five homologous developmental regulation transcription factors: TFAP2A–E, and each transcription factor is coded by independent genes. In structure, the TFAP2 protein contains a highly conserved C-end helix–span–helix sequence motif, the basic DNA-binding domain, a third less conserved domain facing the N end, and the activation domain containing large amounts of proline and glutamine. These factors have been proven to be combined with the GC-rich DNA recognition sequence in a palindromic sequence to become a homodimer or heterodimer, and the promoter was used in specific modes as the transcriptional activator or inhibitor.^32^

Previous work showed that TFAP2C played a different role in various tumors. In most reports related to breast cancer, the overexpression of TFAP2 promoted the occurrence and metastasis of tumors, which enhanced the tumor progress.^33,34^ However, it has also been reported that the decline in TFAP2α expression indicated increased recurrence risks of breast cancer.^35,36^ Interestingly, according to related reports on melanoma, gastric cancer, and colorectal cancer, TFAP2 was an effective cancer suppressor gene.^37–41^ In the highly metastatic A375SM melanoma cells, re-expression of TFAP2 can reduce their oncogenicity and inhibit their metastasis potential in nude mice. Similar research showed that in melanoma, MMP-2 imbalance can be caused by inhibiting TFAP2, which would not promote the oncogenicity.^42^ Another outstanding study showed that miRNA214 can coordinate the formation of metastasis in the melanoma progress model, by directly acting on multiple surface proteins such as TFAP2C and ITGA3.^43^ According to a study of 481 gastric cancer samples conducted at Sun Yat-Sen University, it was found that low TFAP2 expression can be used to independently predict overall poor survival rates of gastric cancer patients.^44^

Previous research proved that the tumor inhibition activity of TFAP2 was mediated through direct interaction with TP53. In addition, TFAP2α can induce TP53-dependent p21 transcriptional activation^45^. On the contrary, a study on a breast cancer cell line showed that no matter what the TP53 state was, the down-regulation of TFAP2α definitely reduced the chemosensitivity.^46^ We found that when TE-1/DDP/exo worked on TE-1 or there was overexpression of miRNA193, it caused silence of TFAP2C and reduced TE-1’s sensitivity to cisplatin. Through positive regulation of p21, TFAP2 mediated its role as differentiation-related transcription factor to negatively regulate the cell cycle. The p21 promoter contained TFAP2α binding sites located at loci −103 and −95, with which TFAP2α was directly combined to stimulate its expression.^46,47^ Furthermore, related research showed that TFAP2α can induce the p21-dependent expression of P53, which proved the potential of TFAP2α in inducing the cell cycle arrest in G1 and G2.^48–50^ After TE-1 obtaining the exogenous exosome TE-1/DDP/exo or the overexpression of miRNA193, the cell cycle of TE-1-Exo or TE-1/miRNA193 mimic presented significant changes, and cell cycle arrest was removed. Based on our research, we believe that the main cause is due to the effects of exosome or overexpressed miRNA193 and the target gene TFAP2C.

## Disclosure

### Ethics approval and informed consent

The experiment was approved by the Ethics Committee of Medical School, Zhengzhou University. All experimental procedures and animal care were approved by the Experiment Center, School of Basic Medical Sciences, Zhengzhou University.

### Consent for publication

All authors agree to publish.

### Availability of data and material

All data generated or analysed during this study are included in this published article (and its supplementary information files).

### Authors’ contributions

XYZ and QXZ planned and designed the research. SFS finished the experiments. XH performed the data analysis. XM prepared the reagents used in the experiments. SFS wrote the manuscript. All authors have read and approved the manuscript.

## Acknowledgements

Not applicable.

## Notes

### Abbreviations

AGE: agarose gel electrophoresis
bp: base pairs
DDP: cis-diaminedichloroplatinum
EV: extracellular vesicles
GO: gene ontology
IC50: the median inhibitory concentration
miRNA: microRNA
nt: nucleotide
PCA: principal component analysis
qPCR: quantitative real-time polymerase chain reaction
RIN: RNA integrity number
TPM: per million

